## Supplemental Figures for "Sensory readout accounts for adaptation"

**Supplemental Materials**

**Supplemental Figure 1**. Response model. Encoding of stimulus is assumed to be a noisy process whereby the distribution of encoded orientations is described by a Gaussian pdf with mean μ and standard deviation σ. Dashed line is pdf and solid line is the cdf of encoding distribution. Note that participants are reporting the probes orientation relative to the stimulus so more frequent CCW responses would correspond to a CW perceptual bias. **A:** Example estimation curve with no bias and a very small σ. If the difficulty was set to 𝛿θ=6° (3 sd) than this participant would get essentially all (99.7%) trials correct. **B:** Estimation curve with a μ=-10, this participant would respond CW on almost every trial. **C-D:** Realistic encoding curves. To aid with fitting and to best describe responses, a constant guess rate of 25% was included in the response model fit to participant responses. C: An unbiased distribution with two theoretical stimuli on which the participant responded CW. The left response 𝛿θ=-6° is incorrect. D: A CCW biased distribution results in a higher likelihood for all CW responses.

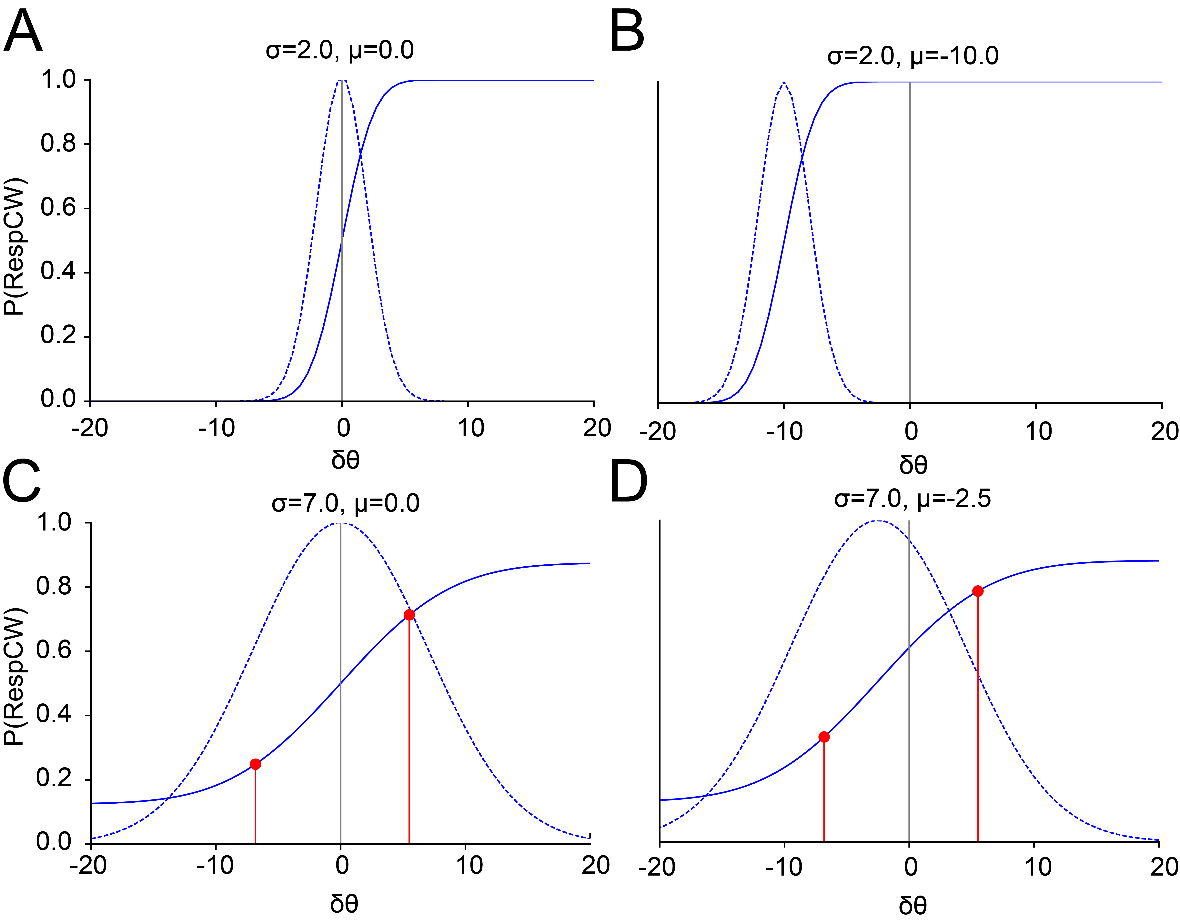

**Supplemental Figure 2.** A subset of participants completed a version of the experiment with inhomogeneities in their stimulus sequences (such that consecutive orientations were not independent). To confirm this manipulation did not drive any of our results, we repeated our behavioral analyses excluding participants with non-independent sequences leaving a cohort of n=25 with an average accuracy of 70.46±1.14° at an average 𝛿θ of 4.97±0.35°. **A,D:** This cohort still showed significant serial dependence (DoG amp =4.71±0.49, *t*(23) = 9.4, p=2.4*10-9; width 0.027±0.0019, FWHM 43.68±1.86°, **B-C:** and had responses that were more accurate (t(24)=3.14, p=.0023, **E-F**: and precise following ‘close’ stimuli (*t*(24)=-3.54, p=0.0009, **G**: Lastly, bias and variance were still positively correlated across this cohort (r(22)=0.72, p=0.00003, **H-J**: Stimulus history effects are larger for worse performing subjects. **H**: Serial dependence was significantly greater for less precise participants (*t*(45)=-2.5, p=.012, unpaired t-test comparing DoG Amplitude). **I-J**: Variance was modulated significantly by stimulus history (low-performing: *t*(23)=3.9 p=.0007; high-performing *t*(22)=2.4, p=.02, one-sample t-tests), with a significant interaction between overall performance and the effect size (p=.017, mixed effects linear model).

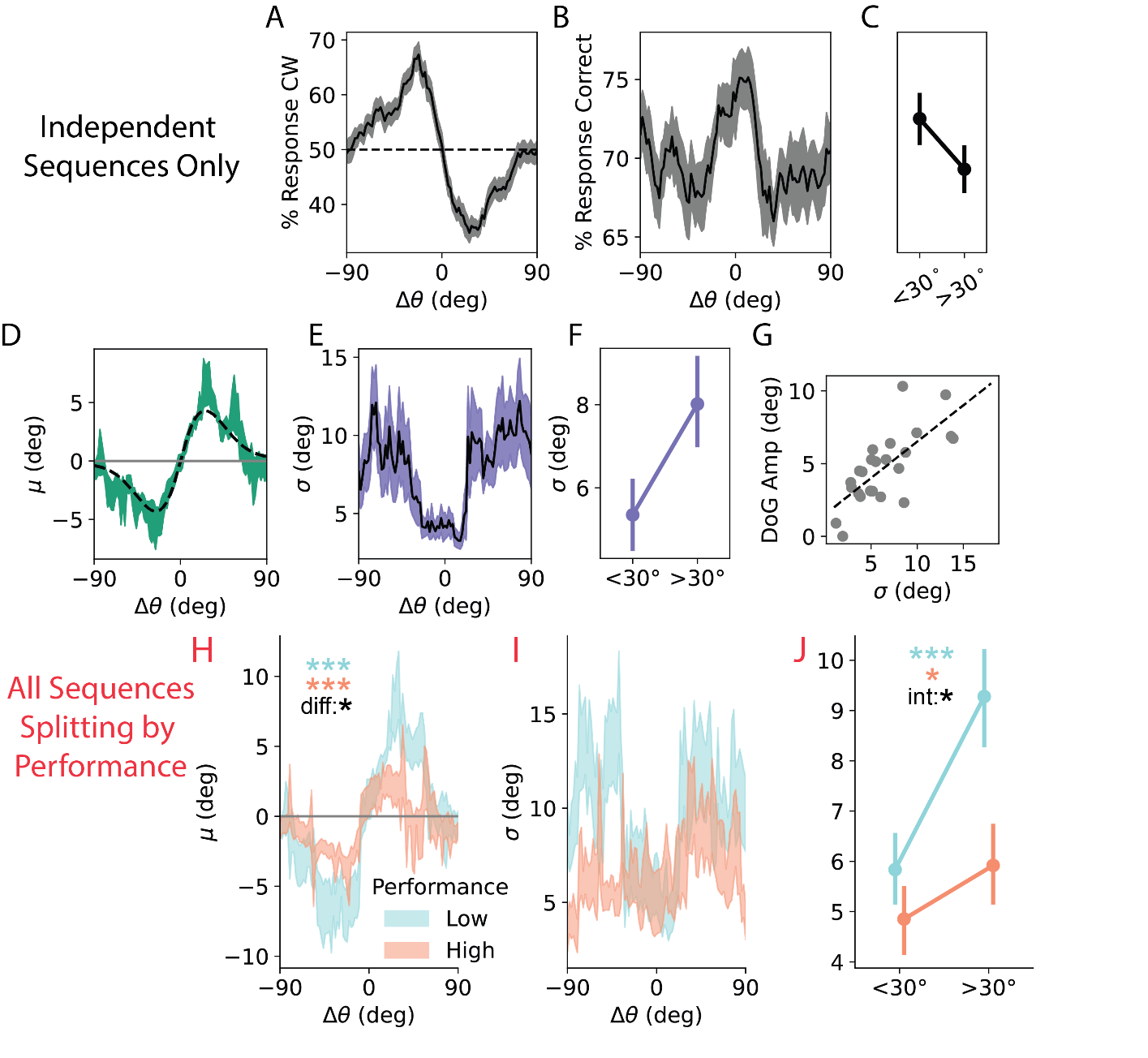

**Supplemental Figure 3.** A subset of these participants completed some sessions where consecutive stimuli were not strictly independent. **A**: To confirm this structure was not driving our results, we repeated the above analysis excluding these sessions and found that responses were still strongly attracted to the previous stimulus (DoG Amp: 3.25± 0.34, *t*(5)=8.85, p=1.53e-04; DoG FWHM: 36.1±2.9). **B**: We found that responses were no longer significantly more precise following small changes in orientation but were trending in the same direction as when including all sessions (*t*(5)=-1.55, p=.09). We additionally confirmed that our finding of reduced bias around small changes in orientation was not driven by the oblique effect in the same manner as the behavioral cohort (mean % cardinal close: 48.6±0.9%, far: 49.8±0.2%, t(5)=-1.0, p=0.36, paired t-test). **C-E**: We further replicate our finding of neural repulsion and increased uncertainty following ‘close’ stimuli across all ROIs except IPS0. **F**: As a control analysis, we attempted but were unable to decode the identity of the next trial in any ROI. ns, not significant; *, p<.05; **, p<.01; ***, p<.001.

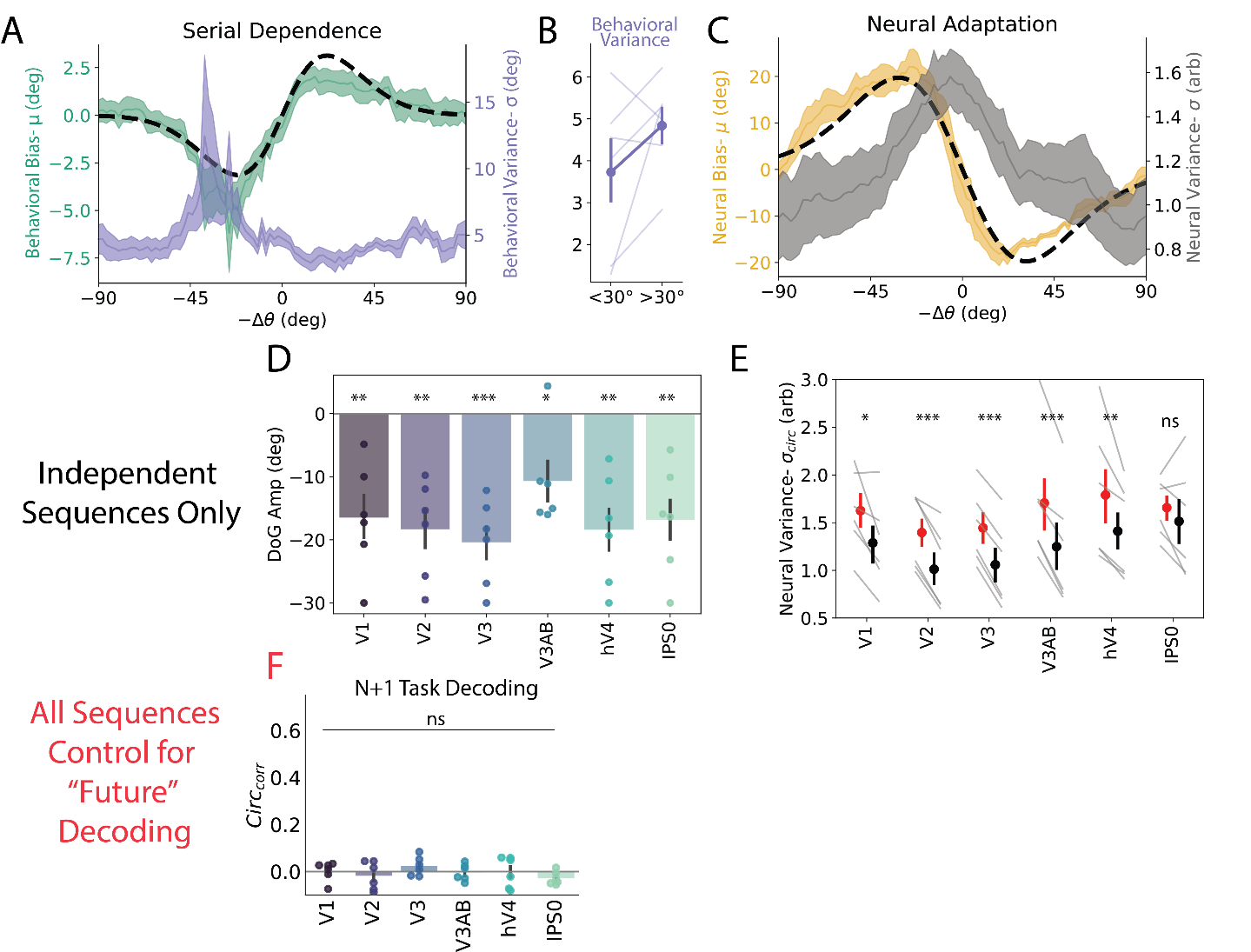

Supplemental Figure 4

Impact of previous trial across time and individuals. **A:** Decoding of the previous stimulus dropped to chance around stimulus presentation before rebounding **B:** decoding using sensory localizer data was consistently at chance during N+1 trial suggesting information relating to past stimulus is not stored in a sensory code. **C-D:** Decoded biases across time for both decoders are consistently repulsive. **E:** Bias curves for individual participants using the memory decoder across rois (see legend) overlayed with behavioral biases (black). Neural and behavioral biases are consistently in opposite directions. Note that id#3 exhibits peripheral repulsion.

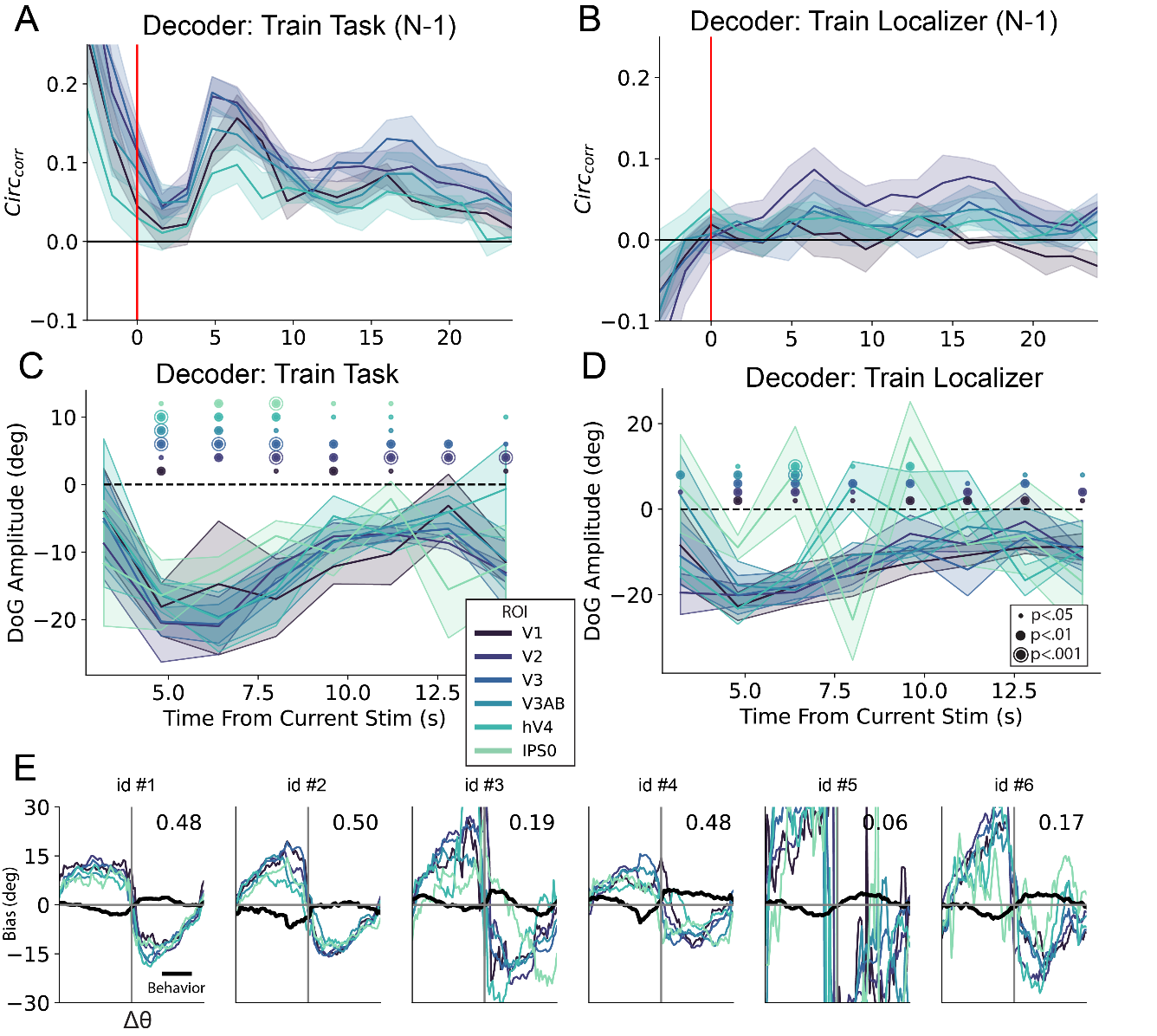

**Supplementary Figure 5.** To quantify the intrinsic dimensionality of neural representations and whether it changes following a ‘close’ stimulus, we performed principal component analyses on the activity matrix (number of trials x number of voxels) of responses across different ROIs. **A:** we found that early principal components were correlated with the presented orientation, here presenting both individual trials as well as the average location for different orientation bins (large solid circles) for an example subject and ROI. **B:** we performed PCA separately for trials following ‘close’ and ‘far’ trials, being careful to subsample the number of trials in the larger group. We then sorted the eigenvalues and examined the proportion of variance explained as a function of the number of components included separately for each group. **C:** we found that it took significantly more components to explain 90% of the variance on the population activity following close versus far stimuli. This suggests that the representations in most visual areas occupy a higher dimensional space following close stimuli, but curiously not V1. Note that the total number of dimensions is shaped by the number of voxels included, so differences between subjects/ROIs should not be interpreted with how these data were processed. **D:** we additionally looked at the area under the variance curve to avoid any arbitrary effects of choosing 90% and found a similar effect (higher AUC implies lower dimensionality).

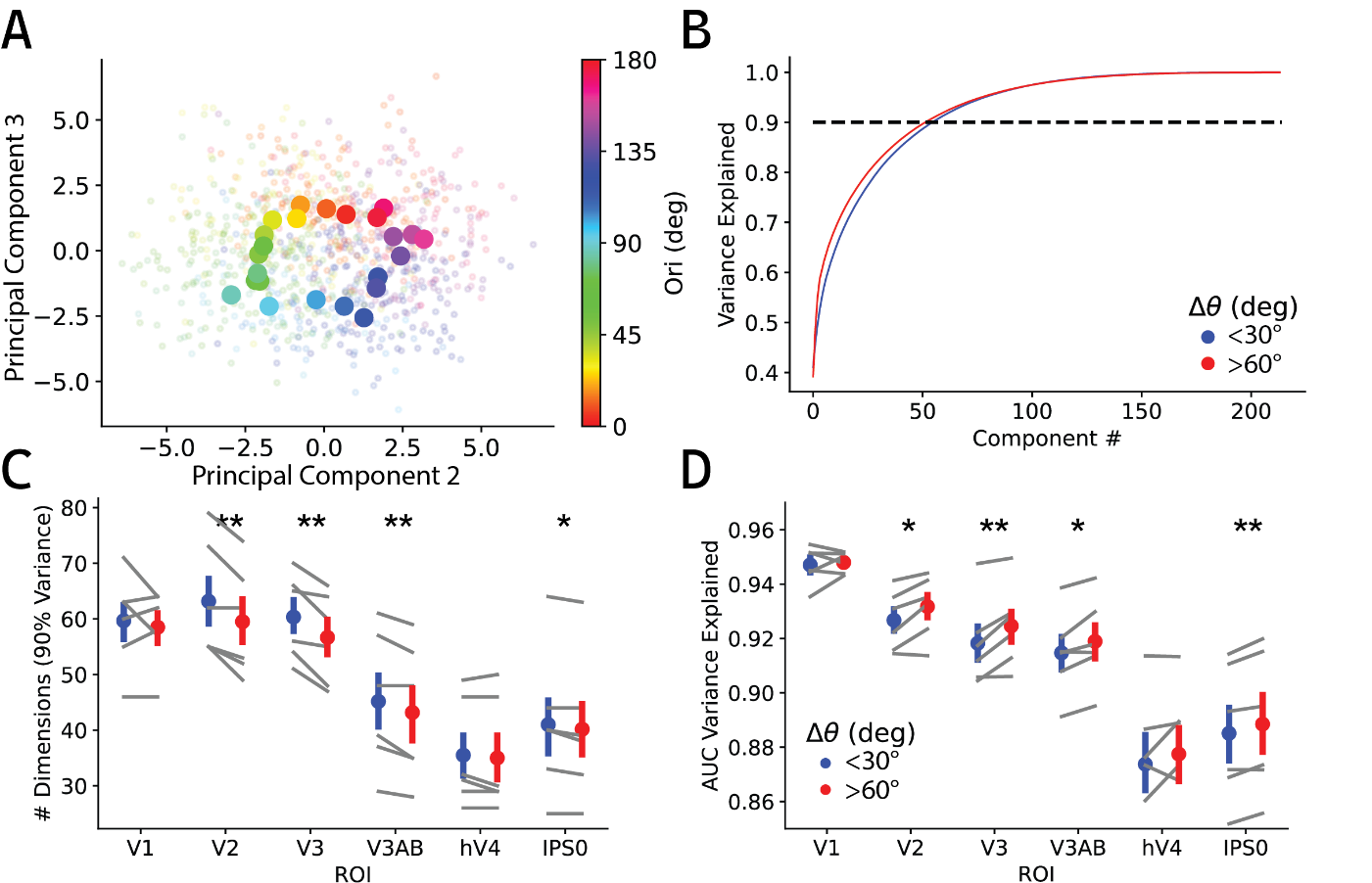

**Supplemental Figure 6.** *Decoded uncertainty as a function of* Δθ across ROIs. **A:** σ_circ_ of decoding errors is significantly greater for close (<30°) versus far (>30°) stimuli across early visual ROIs (see **Neural Variance**). Points and error bars are mean ±SEM across participants; gray lines depict individual participants. Error bars depict SEM across participants **B:** Sliding σ_circ_ for V1-V3 shows a monotonic relationship (± SEM across participants). **C-D:** Same as A-B but measuring uncertainty directly measured from the single trial posterior (see **eq.** **8**). Results are qualitatively very similar for both techniques. *, p<.05, **, p<.01, ***, p<.001.

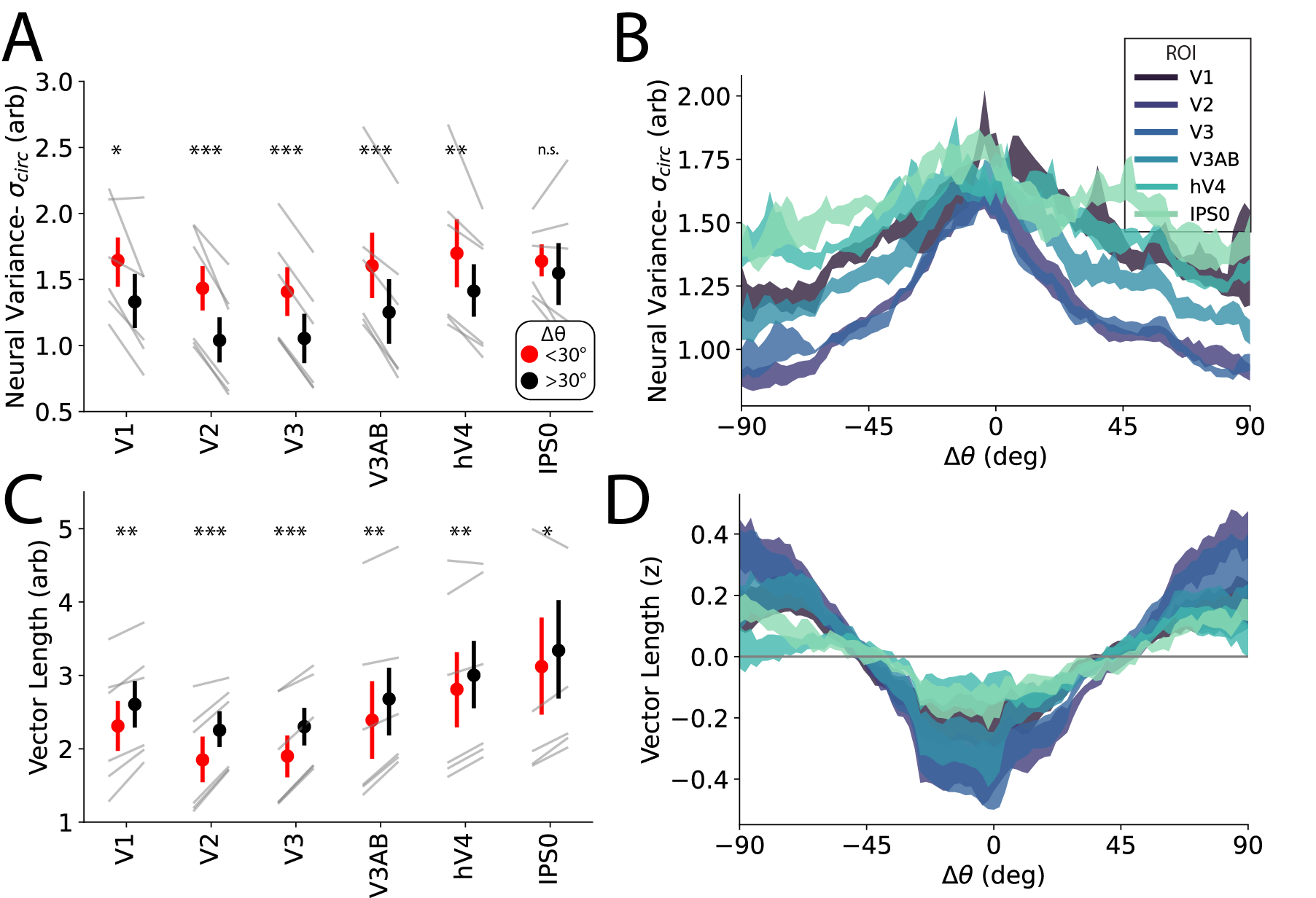

**Supplementary Figure 7.**

To better understand how our experiment’s trial sequence could impact results, we simulated BOLD signals based on our empirically estimated HRFs and our trial sequences used in the task. We first created a population of 32 voxels with uniformly distributed von Mises tuning curves. Note that for the purposes of this simulation, we are effectively treating voxels as neurons instead of a summation of the metabolic demands of many neurons. This shortcut comes from experience simulating voxel activity and finding decoding results are unaffected by such a shortcut while making results a bit simpler to understand (and faster to generate). The responses of each voxel were estimated by first generating a design vector based on the stimulus presentation times of both the stimulus and probe for a given subject with the amplitude of the response based on the defined tuning curves. This vector was then convolved with an empirically estimated HRF (both the raw output and when parameterized with a double gamma function) randomly sampled from voxels of the same subject to get the estimated evoked response to both the stimulus and the probe. These two signals were then combined along with gaussian noise to simulate the voxel response (**A**).

Importantly, the tuning properties of these simulated voxels were unaffected by past stimuli so any biases found by applying our decoding techniques could reflect artifacts of our task design or analysis procedure. We additionally simulated BOLD responses with true adaptation in the underlying neural tuning. For simplicity we simply attenuated the response to the current trial by 40% of the response to the previous trial while keeping all other stages of our analysis the same.

We first applied a decoder across time to the epoched data and found a similar pattern to our empirical data with decoding performance following a parabolic shape before leveling off at some intermediate level, here utilizing HRFs from V3 voxels (**B**). This was true whether we used parameterized or raw HRFs and whether the simulation included adaptation. We next examined biases in our decoder as a function of stimulus history. With adaptation (red curves), decoded representation were systematically repelled from previous stimuli matching our empirical findings (**C**). Importantly, without adaptation the resulting bias was never repelled from the previous stimulus (blue curves). This suggests that the timing of our stimuli and the resulting evoked responses should not bias us towards seeing the repulsive results we report.

We finally implemented the regression-based estimation of BOLD responses as we did with our empirical data. As stated before, this technique should remove any linear contributions of past evoked responses to our estimate of the current trial’s response. When analyzing the resulting biases, we found that while the unadapted data showed no bias from the previous stimulus (as expected, despite added noise) the adapted response continued to show a repulsive bias (**D**).

This analysis demonstrates that 1) while our task design could lead to biases in decoded representations in the absence of any neural history effects, these effects tend to be in the opposite direction of our reported effects and 2) our use of HRF kernels to estimate trial responses is unbiased by across trial contamination and robustly recovers repulsive patterns in the presence of real neuronal adaptation at noise levels similar to our study.

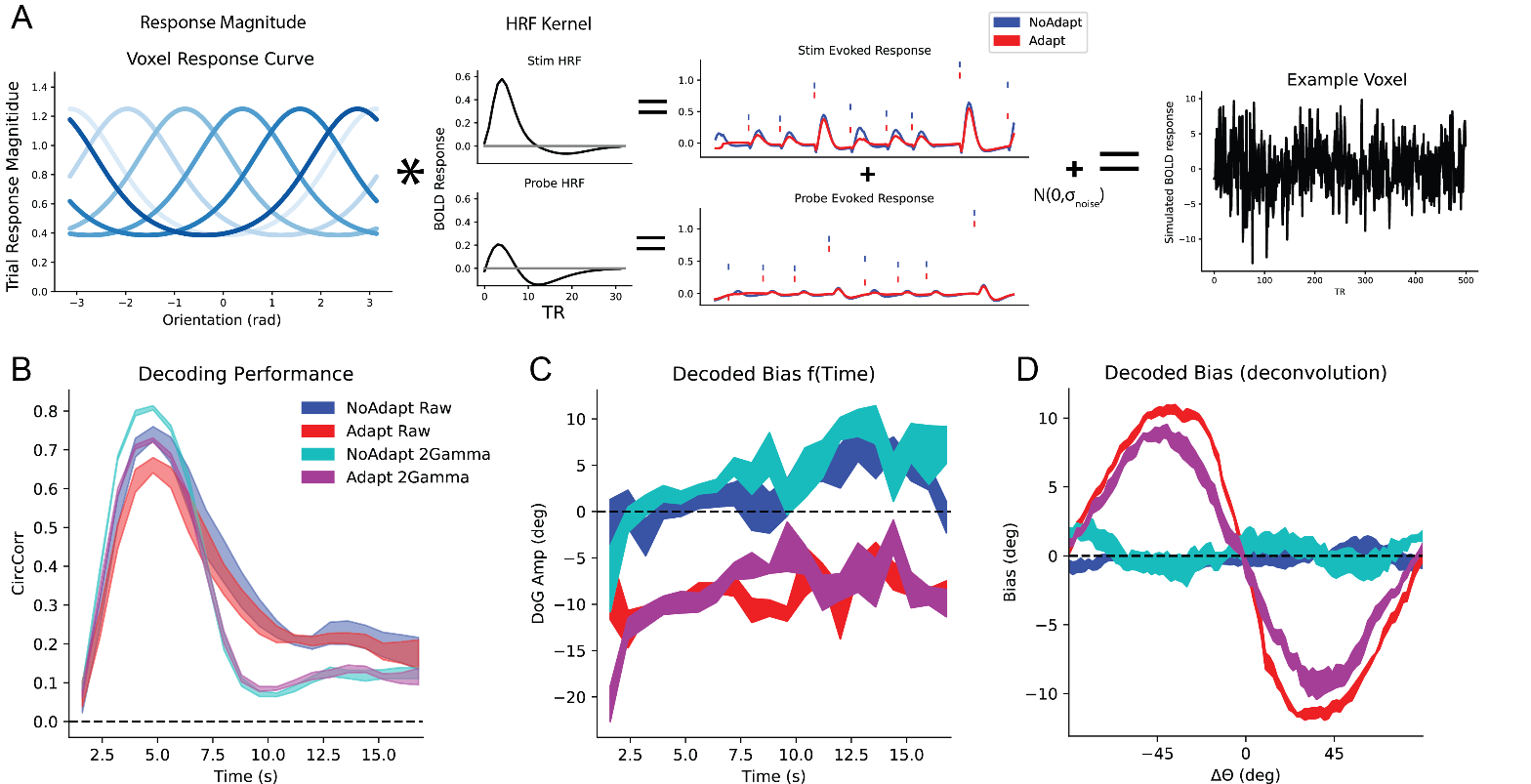

**Supplementary Figure 8.** Model fits for individual participants (same order as Figure 3). Solid lines correspond to empirical neural (yellow) or behavioral (green) bias; dashed lines correspond to model fits to BOLD decoding bias (Unaware model, **A**) or behavior (**B-D**). Model fits plotted are average of noiseless biases generated by models fit to each CV fold. Note that models are fit to raw data, not binned data presented here. Pearson’s correlations are reported above each fit between binned and model estimated bias.

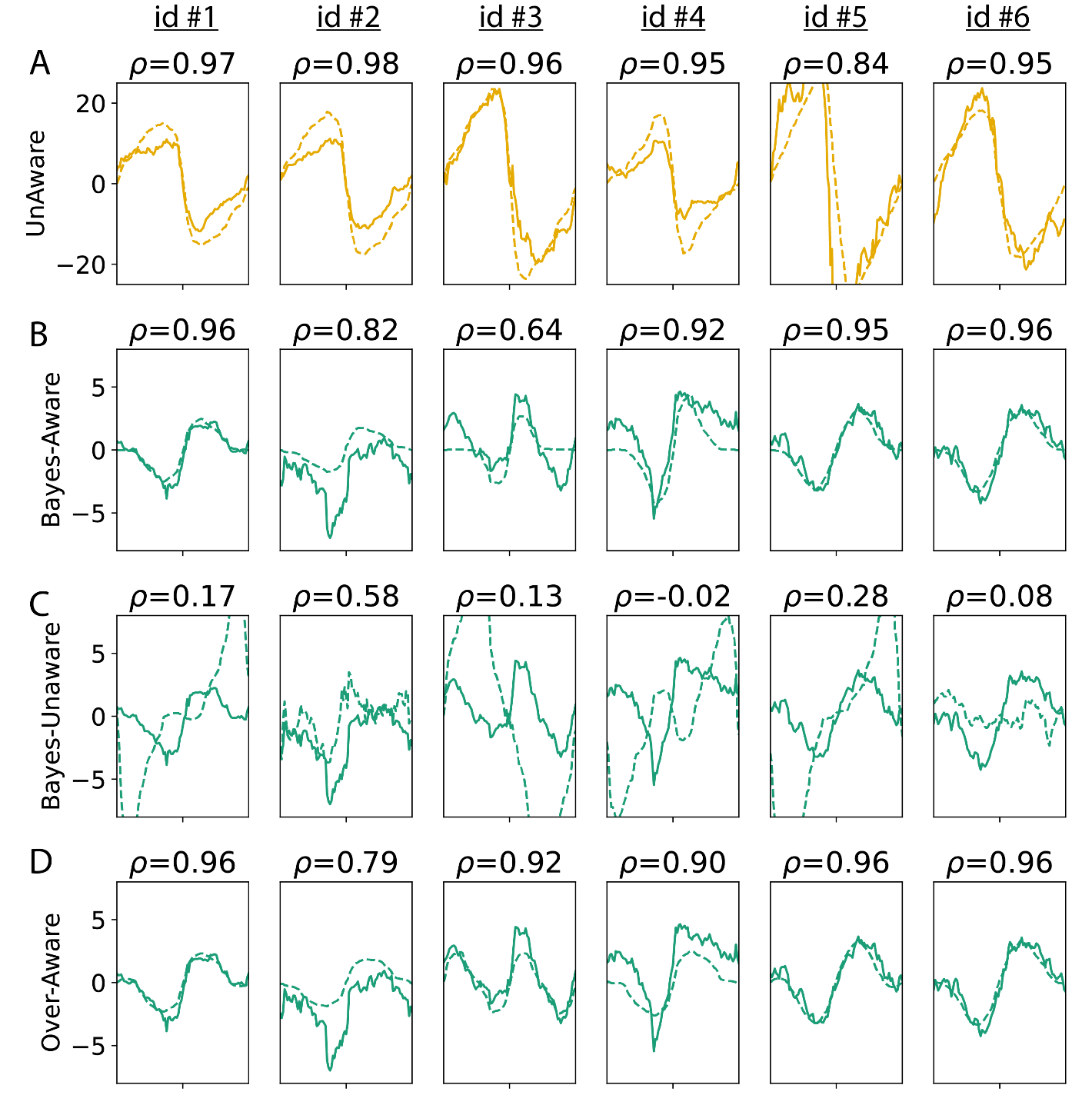

**Supplementary Table 1.** Cells correspond to parameters for proposed decoder. Items with **bold values** indicate free parameters adjusted to fit empirical data (± SEM across participants). $\gamma_{m}$ controls the amplitude and $\gamma_{s}$ controls the width of gain adaptation (Figure 7A). These parameters were fit by minimizing the residual sum of squared errors between the unaware decoder and the BOLD decoder output. $\gamma_{m2} and \gamma_{s2}$ are the assumed adaptation parameters at decoding. These terms were either set to assume no adaptation (unaware), match the true amount of adaptation (aware) or are free parameters adjusted to maximize the likelihood of responses (over-aware, Figure 7B). Last, R adjusts the average Poisson firing rate and ψ controls the variance of the prior distribution (Figure 7C). These parameters are adjusted for decoders using a Bayesian prior while R is set to the arbitrary value of 5 for non-Bayesian decoders (it has no effect on bias for non-Bayesian decoders). Increasing R increases the precision of the likelihood function and reduces the relative influence of the prior. Increasing ψ increases the range of Δθ over which the prior has an influence.

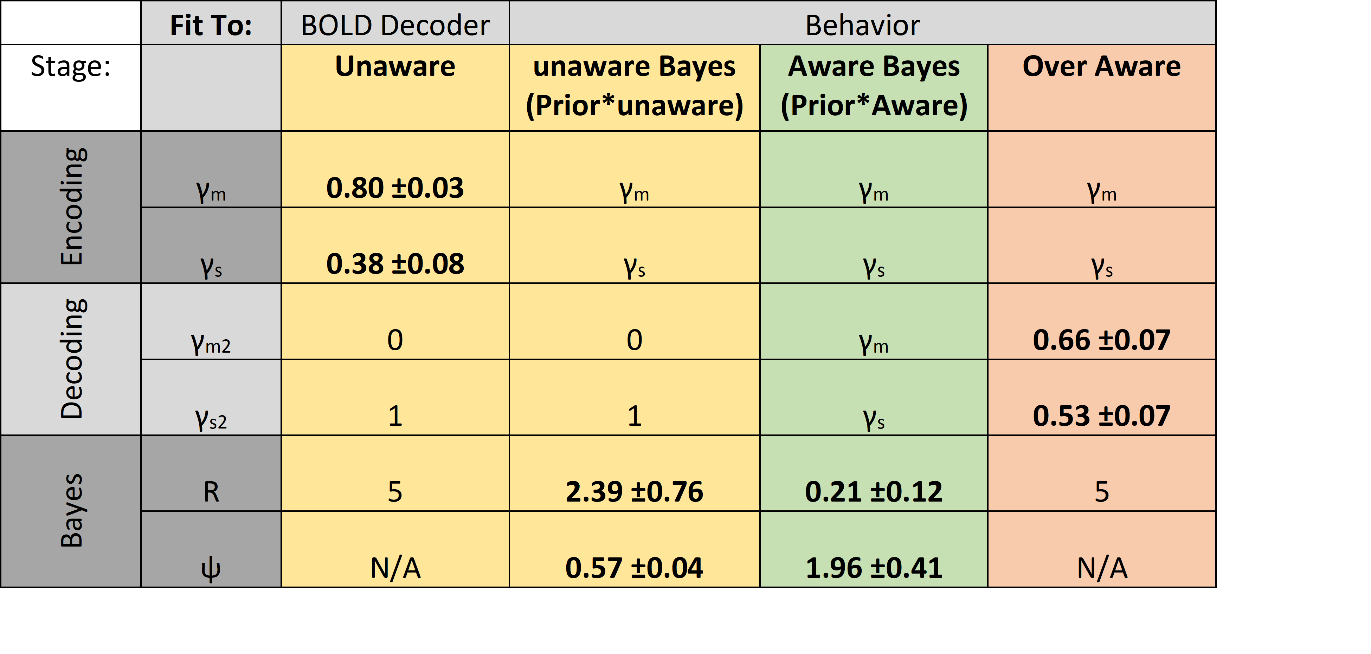
